# DDX17 is required for efficient DSB repair at DNA:RNA hybrid deficient loci

**DOI:** 10.1101/2021.10.14.464298

**Authors:** Aldo S. Bader, Janna Luessing, Ben R Hawley, George L Skalka, Wei-Ting Lu, Noel F Lowndes, Martin Bushell

**Affiliations:** Cancer Research UK Beatson Institute, Glasgow G61 1BD, UK; Centre for Chromosome Biology, School of Natural Science, National University of Ireland Galway, Galway, Ireland, H91W2TY; Department of Pharmacology, Weill Cornell Medicine, Cornell University, New York, NY 10065, USA; The Francis Crick Institute, London, NW1 1AT, UK; Institute of Cancer Sciences, University of Glasgow, Glasgow, G61 1QH, UK

## Abstract

Proteins with RNA-binding activity are increasingly being implicated in DNA damage responses (DDR). Additionally, DNA:RNA-hybrids are rapidly generated around DNA double-strand breaks (DSBs), and are essential for effective repair. Here, using a meta-analysis of proteomic data, we identify novel DNA repair proteins and characterise a novel role for DDX17 in DNA repair. We found DDX17 to be required for both cell survival and DNA repair in response to numerous agents that induce DSBs. Analysis of DSB repair factor recruitment to damage sites suggested a role for DDX17 early in the DSB ubiquitin cascade. Genome-wide mapping of R-loops revealed that while DDX17 promotes the formation of DNA:RNA-hybrids around DSB sites, this role is specific to loci that are naturally deficient for DNA:RNA-hybrids. We propose that DDX17 facilitates DSB repair at loci that are inefficient at forming DNA:RNA-hybrids by catalysing the formation of DSB-induced hybrids, thereby allowing propagation of the damage response.

## INTRODUCTION

Accurate repair of DNA damage prevents the mutations that are the driving force behind carcinogenesis. To maintain fidelity our cells have evolved a network of repair pathways that detect and resolve the various forms of DNA damage including: nucleotide adducts, inter-strand crosslinks, collapsed replication forks and strand breaks(1). Double-strand breaks (DSBs) are considered the most toxic form of DNA damage due to their relatively high propensity for causing mutations(2) and, mutations in multiple DSB repair genes are associated with an increased cancer risk(3, 4).

Initial signalling in response to DSBs is carried out by the three apical DDR kinases: ataxia telangiectasia mutated (ATM), ataxia telangiectasia and rad3-related (ATR) and DNA-dependent protein kinase (DNA-PK)(5). An early and essentially universal marker of DSBs is the phosphorylation of Serine 129 of histone H2AX to form γ-H2AX which directly recruits phosphorylated MDC1 to DNA breaks. The E3-ubiquitin ligase RNF8 then binds to MDC1 and mono-ubiquitylates other factors, including L3MBTL2(6) and possibly the H1 linker histone, thereby creating a marker that recruits the E3 ligase RNF168(7, 8). RNF168 is currently thought to poly-ubiquitylate H1, contributing to decompaction of chromatin, and to mono-ubiquitylate H2A and H2AX at two positions; K13 and K15(8, 9). These H2A ubiquitylations facilitate the recruitment of 53BP1 and BRCA1 that determine pathway choice, either non-homologous end-joining (NHEJ) or homologous recombination (HR), respectively. HR is a high-fidelity replication-based process, whereas NHEJ involves rapid processing and ligation of the broken DNA ends that can be error prone. There is growing evidence that the DDR signalling cascade is more complex as a number of additional factors have been implicated, including additional ubiquitin ligases, such as HUWE1(10), as well as RNA binding proteins, such as DROSHA and WRAP53β(11, 12). Chromatin is not only a barrier to DNA repair but also provides the substrate for post-translational modifications and chromatin remodelling enzymes required for ordered recruitment of both repair factors and the chromatin remodelling enzymes essential for repair of and recovery from DNA damage.

There is currently growing evidence for a role for RNA and RNA binding proteins in the DDR and specifically in DSB repair(13–15). A wide variety of RNA binding proteins have already been characterised as DNA repair genes including the splicing factor THRAP3(16) and the DEAD-box helicase Senataxin^(17)^. In addition, both on-going transcription and RNA itself have been implicated in the damage response, with transcription inhibitors(18–20) and RNase treatments(18, 21) both being shown to significantly impair the damage response. While DNA:RNA-hybrids have been detected around active DSBs and shown to be required for efficient repair(12, 17, 19, 22), more recently, RNA polymerase III has been shown to be recruited to DSBs where it catalyses transcription templated from the 3’ ssDNA overhang produced upon resection(15). In addition, many RNA binding proteins have been identified that promote their formation and resolution around break sites(12, 17, 23, 24). Defining the roles of RNA and RNA binding proteins in the DDR is an emerging area requiring characterisation.

DDX17 is another DEAD-box helicase known to exist as two isoforms known, p72 and p82, that result from an alternative translation start site. DDX17 has RNA helicase, annealing and branch migration activity allowing it to remodel complex RNA structures(25–29) and is known to function in microRNA biogenesis(25, 27, 30, 31), ribosomal biogenesis(26, 27, 30) and splicing(26, 27). DDX17 interacts with the miRNA processors DROSHA and DGCR8 and remodels the 3’ flanking regions of pri-miRNAs to enhance miRNA processing. In mice, *Ddx17^−/−^* knockout results in early embryonic lethality due to impaired miRNA and rRNA processing(32).

Here, we use a meta-analysis of proteomic screens to identify novel DNA repair proteins and highlight DDX17 as a potential repair factor. Our investigations provide evidence that DDX17 is important for maintaining genome stability via DSB repair, specifically via facilitating RNF168 recruitment and subsequent histone ubiquitylation at DSBs. Consistent with these observations, cells deficient for DDX17 function display impaired DSB repair in both NHEJ and HR pathways resulting in DNA damage accumulating to toxic levels. Additionally, we characterised a requirement for DDX17 in promoting DSB-induced DNA:RNA-hybrid formation. Interestingly, this role is preferentially required around DNA breaks in loci that are naturally deficient for DNA:RNA-hybrids. Our data suggest that DDX17 specifically facilitates DSB repair by promoting DSB-induced hybrid formation at regions of the genome which are inefficient at forming hybrids.

## MATERIAL AND METHODS

### Cell culture

U2OS, A549, DIvA and BJ-5ta cells were cultured in Dulbecco Modified Eagle’s Medium (DMEM, GibCo) supplemented with 10% Fetal bovine serum and 2mM L-glutamine with DIvA cells also containing 1μg/mL puromycin. All cells were incubated at 37°C with a 5% CO_2_, humidified atmosphere. hTERT immortalized RPE-1 (ATCC) cells were cultured in DMEM-F12 media supplemented with 10% FBS (Gibco) and 1% PenStep. hTERT-RPE1 cells were verified by STR analysis (Eurofins Genomics). U2OS, U2OS-derived (DIvA, EJ5, HR-U2OS) and hTERT-RPE1 cell lines are female, while A549 and BJ-5ta cell lines are male.

### Transfection and Drug treatments

For DIvA, A549 and BJ-5ta cells, Dharmafect 1 was used at a final dilution of 1/1000 and siRNA targeting either DDX17 (Thermo, s2062 for A549 cells, s20623 for all other experiments), RAD54 () or an untargeted control (Horizon Discovery, D-001810-01-05) were used at a final dilution of 20nM. Cells were cultured for 24h prior to transfection and treated 48h after transfection. siRNA and Dharmafect 1 (Horizon Discovery, T-2001-03) were first diluted separately in optimem to a volume of 5% the final desired medium volume, and incubated for 5min at RT. siRNA and Dharmafect 1 were then combined in 1:1 ratio and incubated for 20 min at RT. The remaining 90% of the final volume of medium was then added and the medium on the cells was immediately replaced with this. For siRNA transfections in hTERT-RPE1, U2OS and HR-U2OS, cells were transfected with Oligofectamine Reagent (LifeTechnologies) according to manufacturer’s instructions. Briefly, 1.2-1.5×10^5^ cells were plated on a 35mm cell culture dish. After 24h cells were transfected with 40pmol of negative control siRNA (Dharmacon) or siRNA targeting the gene of interest. Cells were treated and harvested 48h post transfection. ATMi (KU55933, SelleckChem) and DNA-PKi (NU7026) were used at 10uM unless otherwise stated. Olaparib (AZD2281, SelleckChem) and ICRF-193 (I4659, Sigma) were used at indicated doses. Irradiation was performed using a Mainance Millenium Sample Irradiator containing a Cs-137 sealed source.

### Western blotting

Cells were harvested by lysing in 1.2x sample loading buffer and sonicated with a Diagenode Biorupter for 5 minutes on high to shear the DNA. Samples were run on either 6% or 10% polyacrylamide gels and transferred in tris-glycine transfer buffer containing 20% methanol and 0.01% SDS onto nitrocellulose membranes. Membranes were blocked using 5% BSA in TBST and primary antibody probing was done overnight at 4°C with 5% BSA in TBST. Antibodies and dilutions are listed in Supplementary Table 1. Membranes were then washed for 10 min in TBST 3 times at RT and incubated with secondary antibodies (Li-COR Biosciences) at RT for 1 h at a dilution of 1/10,000. Membranes were then washed for 10 min in TBST 3 times at RT and imaged with a Li-COR Odyssey. For U2OS and hTERT-RPE1 cells, cells were lysed in Lysis Buffer (150mM NaCl, 50mM Tris-Hcl, pH7.5, 10% Glycerol, 0.5%NP-40, 1mM MgCl2, 1/1000 Benzonase (Sigma), Phosphatase (PPI) and Protease Inhibitors(PI) for 45min on ice. After pelleting the lysed cells at 14000rpm for 15min at 4°C, total cell extracts (TCE) were collected and the concentration was measured using Bradford Reagent (Sigma). 25ug of TCE was run on a SDS-PAGE gel at 180V. Proteins were transferred onto a nitrocellulose membrane for 70min at 100V on ice in ice-cold transfer buffer. Membranes were blocked with TBS-T (1xTBS + 0.1% Tween-20) containing 5% milk for 10min at RT prior to primary antibody incubation overnight. After secondary AB incubation, lane were detected by chemiluminescence on a Vilbur FX6 imager.

### Clonogenic survival assay

For U2OS or hTERT-RPE1 cell lines, cells were trypsinized 48h after siRNA transfection and counted. 500 cells were plated onto a 60mm dish. For IR sensitivity assays, cells were allowed to adhere for 1h prior to irradiation at indicated doses (Mainance Millenium Sample Irradiator containing a Cs-137 sealed source), while for Olaparib and ICRF-193 treatments, cells were directly plated in media containing the indicated dose for the duration of the experiment. Cells were grown at 37°C for 10-14 days until colonies were an average of 1-2mm in diameter. Colonies were stained with DMMB (0.25% Dimethylmethylene blue in 50% Methanol) and counted.

### Neutral Comet Assay

Cells were treated with siRNA 48h prior to IR (3Gy, Mainance Millenium Sample Irradiator containing a Cs-137 sealed source) and harvested at indicated timepoints. Neutral comet assays were carried out according to manufacturer’s guidelines (Trevigen). Briefly, cells were harvested and combined with LMAgarose (Trevigen) at a final concentration of 1×10^5^ cells/ml and loaded onto polylysine slides. The agarose plugs were allowed to solidify at 4°C for 1h before being immersed in lysis buffer (Trevigen) overnight at 4°C. Slides were then equilibrated in cold Electrophoresis buffer (100mM Tris pH9.0, 300mM NaAz) for 30min prior to electrophoresis for 1h at 24V. The DNA was precipitated for 30min at RT in DNA precipitation buffer (1M NH4Ac in EtOH) and washed with 70%EtOH for a further 30min. Slides were allowed to dry overnight at 37°C prior to staining with SYBR green (Roche, S7563). Images were acquired on a DeltaVision integrated microscope system using the Applied Precision SoftWoRx acquisition software mounted on an IX71 Olympus microscope with a 10x air objective (Imsol). All images were taken as single slices using a CoolSNAP HQ2 ICX-285 CCD camera. Comet analysis was performed using the CometScore software from Tritek Corporation.

### Metaphase spreads

Cells were treated with 2Gy IR (Xstrahl RS320) and allowed to recover for 48h. Cells were then captured in metaphase by treating with 200ng/mL colcemid (Sigma, D7385) for 2h and were then harvested by trypsinisation as well as their growth medium. Cells were pelleted and resuspended in 10mL 75mM potassium chloride and allowed to swell by incubating at 37°C for 30 min. 5mL of ice-cold fixative (75% methanol, 25% acetic acid) was then slowly added. Cells were then twice pelleted, resuspended in 10mL fixative and incubated on ice for 2 min. Cells were pelleted and resuspended in 4mL of fixative, then dropped onto glass slides from a height of 15-20cm using a p200 pipette. Slides were then steamed over a water bath for 10 s and allowed to dry overnight. The slides were then stained by immersing them in water containing 0.1μg/mL DAPI and then washed by re-immersing in water and again allowed to dry overnight before coverslips were mounted with Vectashield anti-fade mounting medium (Vector Laboratories, H-1000). Spreads were imaged using a Carl Zeiss LSM 710 confocal microscope and counted manually in ImageJ.

### Immunofluorescence

Cells were cultured on glass coverslips and either treated with IR, for U2OS, A549 and RPE-1 cells, or with 300nM hydroxytamoxifen (Sigma, H6278) for 4 hours, for DIvA cells. Cells were washed once in PBS and then pre-extracted at RT for 3 min in CSK buffer (100mM NaCl, 300mM sucrose, 3mM MgCl_2_, 10mM PIPES pH 7.0, 50mM NaF, 5mM Sodium orthovanadate, 10mM β-glycerol phosphate and 0.7% Triton X-100). Cells were then washed once in CSK buffer, once in PBS and then fixed using 4% paraformaldehyde in PBS for 20 min at RT. Cells were then washed once in PBS, once in TBST and then blocked with TBST containing 10% goat serum (Merck, G9023). Cells were then washed twice in TBST for 5 min at RT and incubated overnight at 4°C with primary antibodies diluted in TBST containing 1% goat serum. All primary antibodies and dilutions are listed in Supplementary Table 1. Cells were then washed 4 times with TBST for 5 min at RT before incubating with Alexa-Fluor conjugated secondary antibodies at RT for 1h. Cells were then washed 4 times in TBST for 5 min, dipped in water to remove residual buffer and then mounted on glass slides using Vectashield anti-fade hard-set mounting medium containing DAPI (Vector Laboratories, H-1500). Slides were imaged using a Carl Zeiss LSM 710 confocal microscope and images were analysed using the Fiji distribution of ImageJ(33) via the FindFoci plugin(34). FindFoci settings were calibrated on 4 randomly selected images and these settings were then used to count the foci in all imaged nuclei. For U2OS and hTERT RPE1 WT cells, cells were transfected 48h prior to treatment. Cells were then fixed with 4%PFA (EMS) for 10min at RT before permeabilisation with 0.25% Triton-X100 in PBS for 2min at RT. After blocking with 1%BSA in PBS for 1h, cells were incubated for 1h with primary antibody at 37°C and subsequent secondary antibody for 1h at 37°C. Slides were mounted using Vectashield mounting media with DAPI (VectorLaboratories, H1200). Images were acquired on a DeltaVision integrated microscope system using the Applied Precision SoftWoRx acquisition software mounted on an IX71 Olympus microscope with a UPLFLN 40x objective (numerical aperture [NA] 1.3) (Imsol). All images were taken as Z-slices (0.5μm thickness) using a CoolSNAP HQ2 ICX-285 CCD camera. Parameters for image acquisition were kept constant throughout each experiment. Images were deconvolved using SoftWoRx conservative deconvolution. Quantification was carried out using FIJI software(33). In brief, after deconvolution, the images were projected using ‘sum slices’ (FIJI software) to prevent data loss, and pixel clusters above a defined threshold were counted as individual foci. For representative images, the slices were projected using ‘max projection’.

### DSB reporter assays

For the DR-GFP assays, HR-U2OS cells (Luessing et al 2021, under review) were used, while for the NHEJ (EJ5) U2OS derivative cell line was kindly provided by J. Stark^76^. 2×10^6^ cells were transfected with 5ug pCBA-I-SceI plasmid (Addgene #26477), 40nmol siRNA and 1ug Cerulean-c1 plasmid (Addgene #54604) using electroporation (BioRad electroporator). Cells were harvested and resuspended in 500ul PBS containing 40nM TOPRO-3 iodide (Life Technologies, #T3605) to identify live cells. The cells were gated for live cells, doublet exclusion and transfected cells (cerulean-positive). A minimum of 20.000 transfected cells were then assessed for the expression of GFP. FACS analysis was carried out using BD FACSCANTOII and BD-FACS DIVA software. The remaining cells were used for checking the knock-down efficiency by Western Blotting.

### DNA:RNA-immunoprecipitation (DRIP)

DIvA cells were seeded on 15cm plates for 24 h, transfected for 48 h and then treated with 300nM hydroxytamoxifen for 4 h before harvesting with trypsin. Cells were lysed in cytoplasmic lysis buffer (50mM HEPES pH7.9, 10mM KCl_2_, 1.5mM MgCl_2_, 0.34M sucrose, 0.5% triton, 10% glycerol, 1mM DTT) for 10 min on ice and then washed once in cytoplasmic lysis buffer. Nuclei were pelleted at 1500g for 5 min at 4°C and then resuspended in genomic extraction buffer (50mM Tris pH 8.0, 5mM EDTA pH 8.0, 1% SDS and 0.5mg/mL Proteinase K (Invitrogen, 25530049)) and incubated at 55°C for 1h to digest all proteins. The genomic DNA was then precipitated by adding 0.1 volumes of 3M Sodium acetate pH 5.2 and then adding 2.5 volumes of 100% ethanol and incubating at RT until the DNA visibly precipitated. The DNA was spooled onto a P1000 pipette tip, washed once in 75% ethanol, air dried and resuspended in 300uL water. 50μg of DNA was diluted into 300uL of water and then sonicated for 10 cycles of 5s on/30s off on the low setting in a Diagenode Biorupter and run on a 1.2% agarose gel to ensure fragmentation. 25ug of fragmented DNA was then equilibrated to 1X RNase-H buffer and treated with 50 units of RNase-H (NEB, M0297L) for 3h at 37°C, then topped up with an additional 50 units of RNase-H and incubated for a further 3h. All samples, including RNase-H treated controls, were then equilibrated to 1X DRIP buffer (50mM Tris pH 8.0, 5mM EDTA pH 8.0, 140mM NaCl, 1% Triton X-100) and 20μg was immunoprecipitated overnight at 4°C with 10μg S9.6 antibody (Merck, MABE1095) conjugated to 100μL Pierce ChIP-grade protein-A/G magnetic beads (Thermo, 26162). Beads were then washed once in 1X DRIP buffer, once in 1X DRIP buffer with 500mM NaCl, once in LiCl buffer (10mM Tris pH 8.0, 250mM LiCl, 1mM EDTA pH 8.0, 1% NP-40) and then twice in 1X TE buffer. DNA was eluted from the beads by incubation in 100 μL genomic extraction buffer for 30 min at 55°C. The eluted DNA was then purified via phenol:chloroform:isoamylalcohol (pH 8.0, Sigma P2069) and subsequently ethanol precipitated. The DNA was then either analysed by qPCR using Fast SYBR Green (Thermo, 4385618) or used for a DRIP-seq. qPCR primers used are listed in Supplemental Table 2.

### DRIP-Seq

5ng of DRIP DNA was library prepped using the Ultra II DNA library prep kit for Illumina (NEB, E7645L) according to the manufacturer protocol using 6 PCR cycles. Libraries were then subjected to paired-end 75 cycle sequencing on an Illumina NextSeq 500 an the raw data is available at Array Express, accession number E-MTAB-10814. Analysis was done using custom Bash and R scripts, all scripts are available at https://github.com/Bushell-lab/drip_seq. Briefly, fastq files were aligned using Bowtie2(35) and then sorted and indexed using Samtools(36). Paired alignments were converted to bedpe files using Bedtools and then cut to standard bed files and converted back to bam files. Read coverage was calculated using Samtools Depth and then normalised to total library size. These coverage files were then imported into R for further processing, statistical analysis and plotting.

### Proteomic meta-analysis

Published data from proteomic studies was accessed via publication supplementary materials and filtered for significant hits mostly using the parameters from the original publication. These datasets were compiled into an SQL database using custom python scripts allowing for analysis of hit frequency and dataset overlaps. For protein interaction networks, interaction strengths were generated via the STRING database(37) and were then modelled in Cytoscape(38) with groups being created via statistical clustering using the ClusterONE plugin(39).

## RESULTS

### A meta-analysis of proteomic datasets identifies DDX17 as a potentially novel DNA repair factor

Many large-scale proteomic investigations into the DNA-damage response (DDR) have been conducted and substantial such datasets have been published. Therefore, to identify novel DNA repair factors, we conducted a meta-analysis of 11 such proteomic datasets focussed on DNA repair (Supplemental Fig1. A-B)(40–49). This meta-analysis included 4 major groups of investigations; chromatin association studies(40–43), DNA-repair factor interactomes(44–46), phosphoproteomics(40, 47, 48) and ubiquitylomics(49) which were investigating changes in response to DNA damage (Fig. 1A).

**Figure 1:**
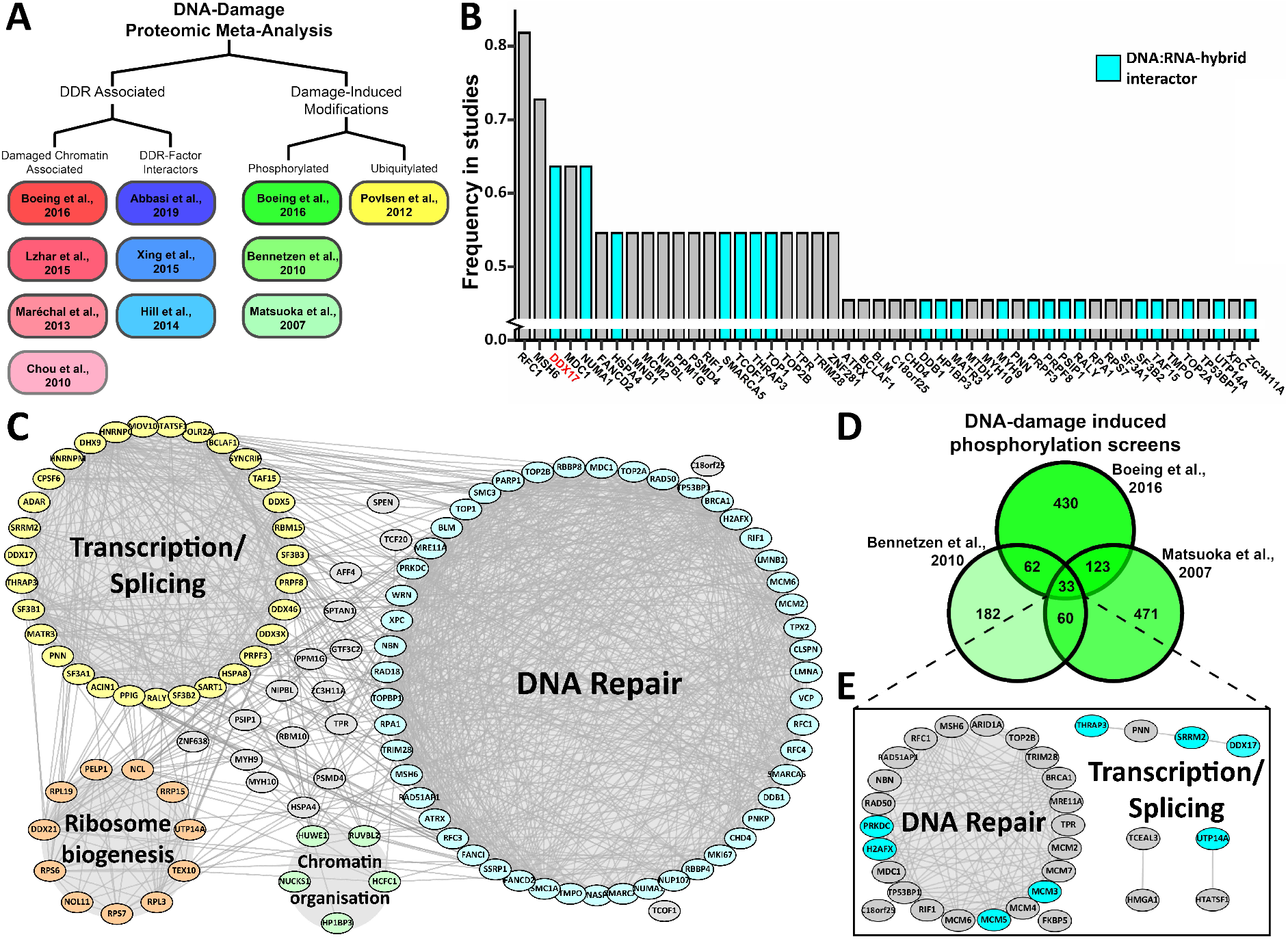
A meta-analysis of proteomic datasets identifies DDX17 as a potentially novel DNA repair factor. (A) Design schematic of the proteomic meta-analysis: sub-groups were designed to ensure a broad range of studies were included in the analysis. (B) Histogram of the frequency in which proteins were identified by the studies in the meta-analysis with DNA:RNA-hybrid interactors highlighted in cyan, only those identified in 5 or more studies are shown. (C) Protein interaction network of all proteins identified in 4 or more studies in the meta-analysis. (D) Venn diagram of the results of the 3 phosphoproteomic studies in the meta-analysis. (E) Protein interaction network of the 33 central proteins from (D).

Protein-protein interaction networking of the 119 proteins identified by over one third of the studies in our meta-analysis revealed 2 major protein clusters; DNA repair and transcription/splicing as well as several other proteins that are partly associated with ribosomal RNA biogenesis and chromatin organisation (Fig. 1C). Gene ontology analysis of the 119 enriched genes confirmed this result, as DNA repair and splicing GO terms were among the most significant along with terms associated with DNA replication and chromosome organisation (Supplementary Figure S1C). Ranking all genes by the frequency in which they were identified in these studies again highlighted commonly identified DNA repair factors, such as MDC1 and MSH6. However, some genes which have not been reported to be DNA repair factors were also significantly enriched, such as DDX17 (Fig. 1B). Interestingly, several of these enriched genes have also been shown to interact with DNA:RNA-hybrids(50), and DDX17 was found to be the most significantly enriched hybrid interactor without a previously described role in DNA repair (Fig. 1B)(51, 52).

A further analysis of the phosphoproteomics studies in the meta-analysis found that only 33 genes are shared between all three studies (Fig. 1D). The low overlap between these studies is likely due to methodological differences, although those identified in all three studies might be expected to be those most likely to be core DNA repair factors and, as expected, a majority of these 33 proteins are indeed canonical DNA repair factors. However, eight were associated with transcription and splicing processes, DDX17 among them (Fig. 1E). Interestingly, all three studies identified the same DDX17 phospho-peptide which harboured 2 “SQ” motifs that are the target of the DDR kinases ATM and ATR (Supplementary Figure S1D). Collectively, this meta-analysis highlighted not only a strong enrichment of transcription and splicing factors in DNA repair processes but, specifically, highlighted DDX17 as a potentially novel DNA repair factor.

### DDX17 is required for cell survival in response to DNA damage via double-strand break repair

To investigate this potential role of DDX17 in the DDR, we initially conducted clonogenic survival assays with RPE-1 cells in response to an increasing dose of irradiation (IR) upon siRNA mediated knockdown of DDX17. In RPE-1 cells, DDX17 knockdown resulted in a significant reduction in cell survival in response to as little as 0.5 Gy IR relative to control cells (Fig. 2A). This reduction in cell survival was comparable to that observed for treatment with the ATM inhibitor KU55933, suggesting an important role for DDX17 in the DDR. IR sensitivity upon DDX17 depletion was also observed in an independent cell line (Supplementary Figure S2A). These U20S cells were also treated with Olaparib and the topoisomerase II inhibitor ICRF-193, agents specific for HR or NHEJ, respectively (Fig. 2B-C). In all of these experiments, DDX17 knockdown was found to significantly reduce cell survival in response to all the DSB-inducing agents used demonstrating a broad role for DDX17 in the DDR. To directly assess the persistence and resolution of DSBs, we next used neutral comet assays. Comet tail moments, the measure of DNA breaks, significantly increased with both control and DDX17 knockdown cells 15 minutes post 3Gy IR. However, 24 hours after irradiation cells in which DDX17 had been depleted had significantly higher comet tail moments than in control cells (Fig.2D-E, Supplementary Figure S2B). This is indicative of a compromised DSB repair and demonstrates that DDX17 depletion significantly reduces DSB repair capacity. To further investigate the direct effect of DDX17 depletion upon maintenance of genome stability, we employed metaphase spreads. DDX17 depleted cells treated with 2 Gy IR and allowed to recover for 2 days (Supplementary Figure S2C) displayed substantial loss of chromosomes (Fig. 2F-G), consistent with sustained, unresolved DNA damage(53, 54). Combined with the comet assay results, this provides evidence for the direct involvement of DDX17 in DSB repair and the maintenance of genome stability.

**Figure 2:**
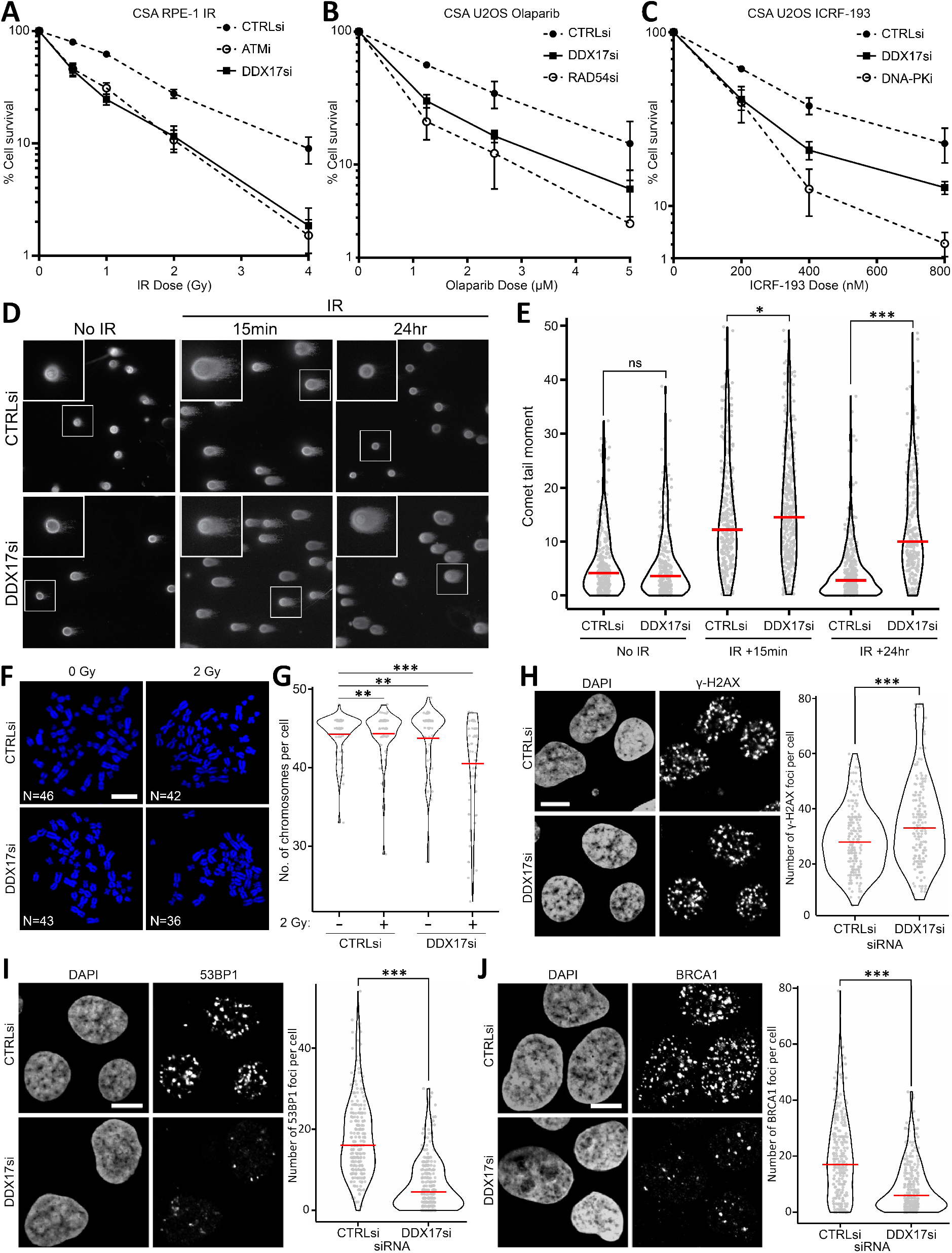
DDX17 is required for cell survival in response to DNA damage via double-strand break repair. (A) Clonogenic assay comparing RPE-1 cell survival in response to increasing doses of IR between the indicated knockdowns and treatments, error bar is SEM of 3 biological replicates. (B) Same as (A) but with increasing Olaparib treatment in U2OS cells. (C) Same as (A) but with increasing ICRF-193 treatment in U2OS cells. (D) Representative images of comet assays with either control and or siRNA with no treatment, 15 minutes post 3Gy IR or 24 hours post 3Gy IR. (E) Comet tail length quantification plot for (D) of a minimum of 268 cells per condition across 2 biological replicates, red line is the median. (F) Representative images of metaphase spreads treated with either control of DDX17 siRNA and either 0Gy IR or 2Gy IR. (G) Number of chromosomes per cell quantification plot for (F) of a minimum of 60 cells per condition across 2 biological replicates, scalebar is 5μM, red line is the mean. (H) Left: representative images of immunofluorescence of γ-H2AX in DIvA cells treated with 4-hydroxytamoxifen with either control or DDX17 siRNA, scalebar is 10μM, right: γ-H2AX foci per cell quantification of a minimum of 250 cells per condition across 3 biological replicates, red line is the median. (I) Same as (H) but for 53BP1 immunofluorescence. (J) Same as (H) but for BRCA1 immunofluorescence. All statistics were done using unpaired, directional Wilcoxon tests, *p<0.05, **p<0.01, ***p<0.001.

Given these pronounced DNA repair phenotypes upon DDX17 knockdown, we quantified the focal recruitment of phosphorylated H2AX (γ-H2AX), 53BP1 and BRCA1 using immunofluorescence. For these experiments, we used the “Damage-Induced via AsiSI (DIvA)” cell system pioneered by the Legube lab that uses an AsiSI restriction enzyme fused to an oestrogen receptor to induce double-strand breaks at known genomic loci in response to 4-hydroxytamoxifen (OHT) treatment. DDX17 depletion resulted in a small but significant increase in yH2AX foci per cell (Fig. 2H), again suggesting that DDX17 knockdown results in an accumulation of unresolved DNA damage. However, immunofluorescence of both 53BP1 and BRCA1 showed substantially reduced focal recruitment of both after DDX17 knockdown (Fig. 2I-J), which cannot be explained by reduced levels of either 53BP1 or BRCA1 upon DDX17 depletion (Supplemental Fig. 2D). We further confirmed these results with IR-induced foci (IRIF) quantifying both γ-H2AX and 53BP1 in U2OS cells. DDX17 depletion once more results in a subtle increase in γ-H2AX IRIF but a significant decrease in 53BP1 foci (Supplementary Figure S2E-G). These results indicate that DDX17 contributes to DDR signalling at an early stage in the cascade. To further quantify the role of DDX17 in DSB repair, we conducted repair assays, using the well-established GFP reporter cell lines DR-GFP and EJ5, which quantify the efficiency of HR and NHEJ, respectively. DDX17 depletion significantly reduced the efficiency of both HR and NHEJ repair efficiency, with the NHEJ reporter efficiency being reduced to a level comparable to 53BP1 knockdown (Supplementary Figure S2H-I). Thus, our data have identified a crucial role for DDX17 in promoting the recruitment of downstream DSB repair factors to facilitate the repair process and ultimately maintain genome stability.

### Immunofluorescence identifies the ubiquitin cascade as the point of DDX17 action in DSB repair

The recruitment of 53BP1 or BRCA1 to DSBs is a relatively early event in the DDR that follows a cascade of histone modifications and protein recruitment. Therefore, we analysed the recruitment of several components of the cascade to determine DDX17’s point of action.

As our previous immunofluorescence experiments found no decrease in γ-H2AX focus formation upon DDX17 depletion in DIvA cells with induced DSBs (Fig. 2H), we next probed for recruitment of the ubiquitin ligase RNF8 which occurs through direct interaction with MDC1. DDX17 depletion did not significantly perturb the formation of RNF8 damage-induced foci (Fig. 3A), suggesting a function downstream of RNF8. Recruitment of the subsequent ubiquitin ligase, RNF168, is dependent upon prior RNF8 recruitment and, interestingly, we observed a significant decrease in RNF168 damage-induced foci upon DDX17 depletion (Fig. 3B). In addition, the abundance of conjugated ubiquitin, detected with a specific antibody, was also similarly reduced upon DDX17 depletion (Fig. 3C). Thus, DDX17 functions downstream of RNF8 to facilitate RNF168 focal recruitment and polyubiquitylation at sites of DNA damage. We used quantitative western blotting of mono-ubiquitylated γ-H2AX, which is dependent upon RNF168 and required for subsequent 53BP1 and BRCA1 recruitment, to support these immunofluorescence data and similarly observed reduced damage-dependent monoubiquitylated γ-H2AX upon DDX17 knockdown (Fig. 3D-E). As there is no change in either RNF8 or RNF168 protein levels (Fig. 3D), the observed defects in recruitment and ubiquitylation result from regulation of protein recruitment to chromatin proximal to DSBs sites, rather than any changes in gene expression. Consistent with RNF168 activity being required for their recruitment, both 53BP1 and BRCA1 display defective focal recruitment to sites of DNA damage (Fig. 3F). We independently validated key results using a different DNA damaging agent and siRNA in A549 cells (Supplemental Fig. 3A-D). Thus, DDX17 is required for the recruitment and activity of RNF168, but not RNF8, to chromatin in the vicinity of DSBs, identifying DDX17 as a crucial factor in the regulation of DSB signalling functioning between these two E3 ubiquitin ligases.

**Figure 3:**
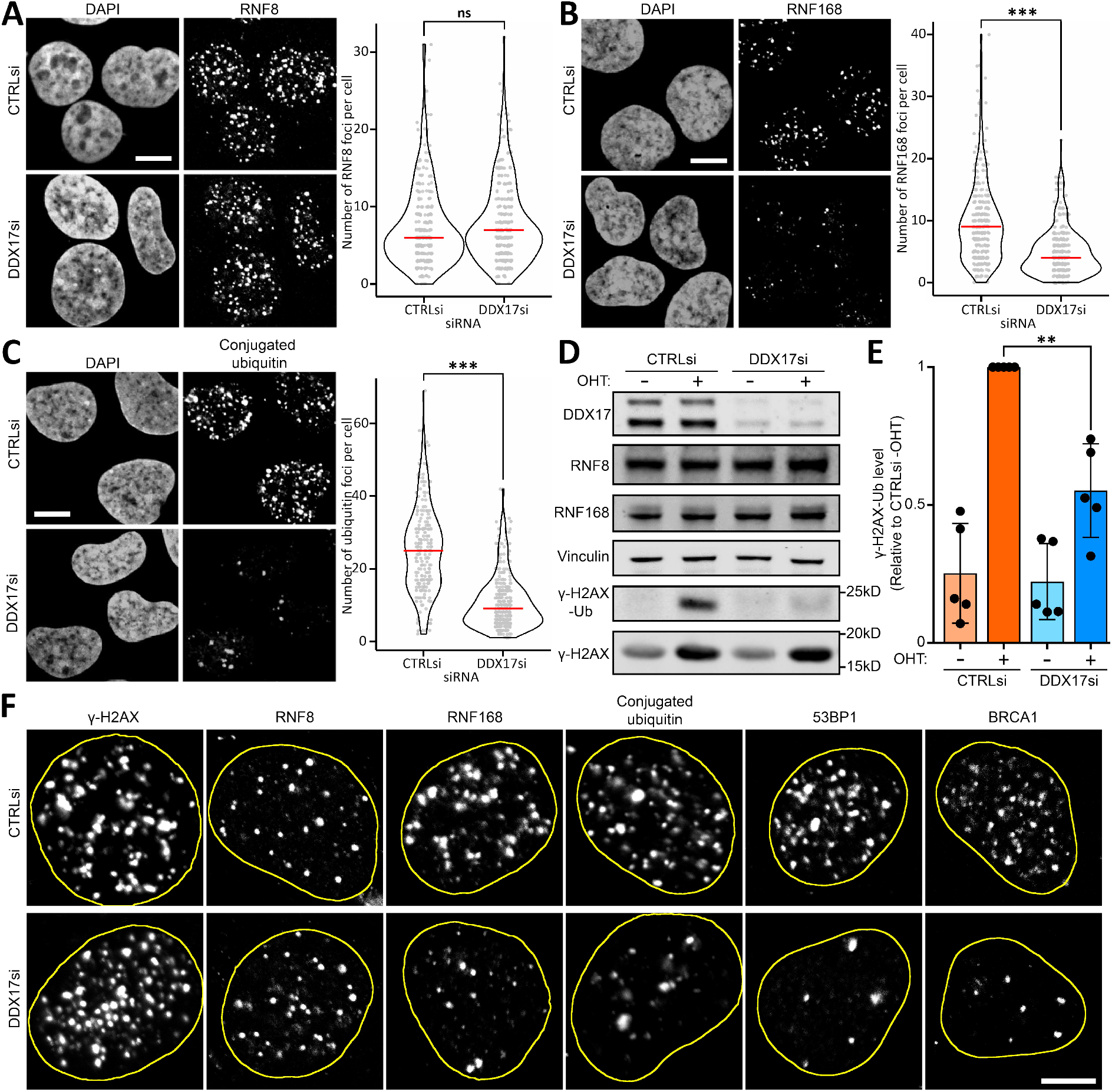
Immunofluorescence identifies the ubiquitin cascade as the point of DDX17 action in DSB repair. (A) Left: representative images of immunofluorescence of RNF8 in DIvA cells treated with 4-hydroxytamoxifen with either control or DDX17 siRNA, scalebar is 10μM, right: RNF8 foci per cell quantification of a minimum of 265 cells per condition across 3 biological replicates, red line is the median. Statistics done using an unpaired directional Wilcoxon test, not significant (ns). (B) Same as (A) but for RNF168 immunofluorescence, ***p<0.001. (C) Same as (A) but for conjugated ubiquitin immunofluorescence, ***p<0.001. (D) Western blots for of DIvA cells transfected with either control or DDX17 siRNA and treated with or without 4-hydroxytamoxifen (OHT), mono-ubiquitylated γ-H2AX is determined by the size shift of the γ-H2AX main band from ∼16kD to ∼24kD. (E) Quantification of γ-H2AX-Ub bands from (D) relative to total γ-H2AX then relative to CTRLsi +OHT. Bar is mean of 5 biological replicates, points are the individual replicate values, error bar is standard deviation. Statistics were done using a paired t-test, **p<0.01. (F) Representative images of immunofluorescence for γ-H2AX, RNF8, RNF168, conjugated ubiquitin, 53BP1 and BRCA1 with either control or DDX17 siRNA treatment, nuclei are outlined in yellow, scalebar is 5μM.

### DDX17 promotes double-strand break induced DNA:RNA-hybrids at hybrid deficient genomic loci

Loss of RNF168 recruitment and reduced ubiquitylation at sites of DNA damage is consistent with the phenotype previously observed for the microRNA biogenesis factor DROSHA(12), a known interactor of DDX17(32, 55). This study found DROSHA to be required for the formation of DNA:RNA-hybrids at DSBs, and that these hybrids were critical for effective DSB repair. Given that DDX17 was also previously reported to be a DNA:RNA-hybrid interactor(12), we chose to investigate the effect of DDX17 knockdown on DSB-induced DNA:RNA hybrid formation. To do this, we used the well-established technique of DNA:RNA-hybrid immunoprecipitation (DRIP) in the DIvA cell system to quantify DNA:RNA-hybrid levels around AsiSI-induced DSBs (Figure 4). Western blotting confirmed the expected induction of g-H2AX and DDX17 depletion (Supplementary Figure S4A).

**Figure 4:**
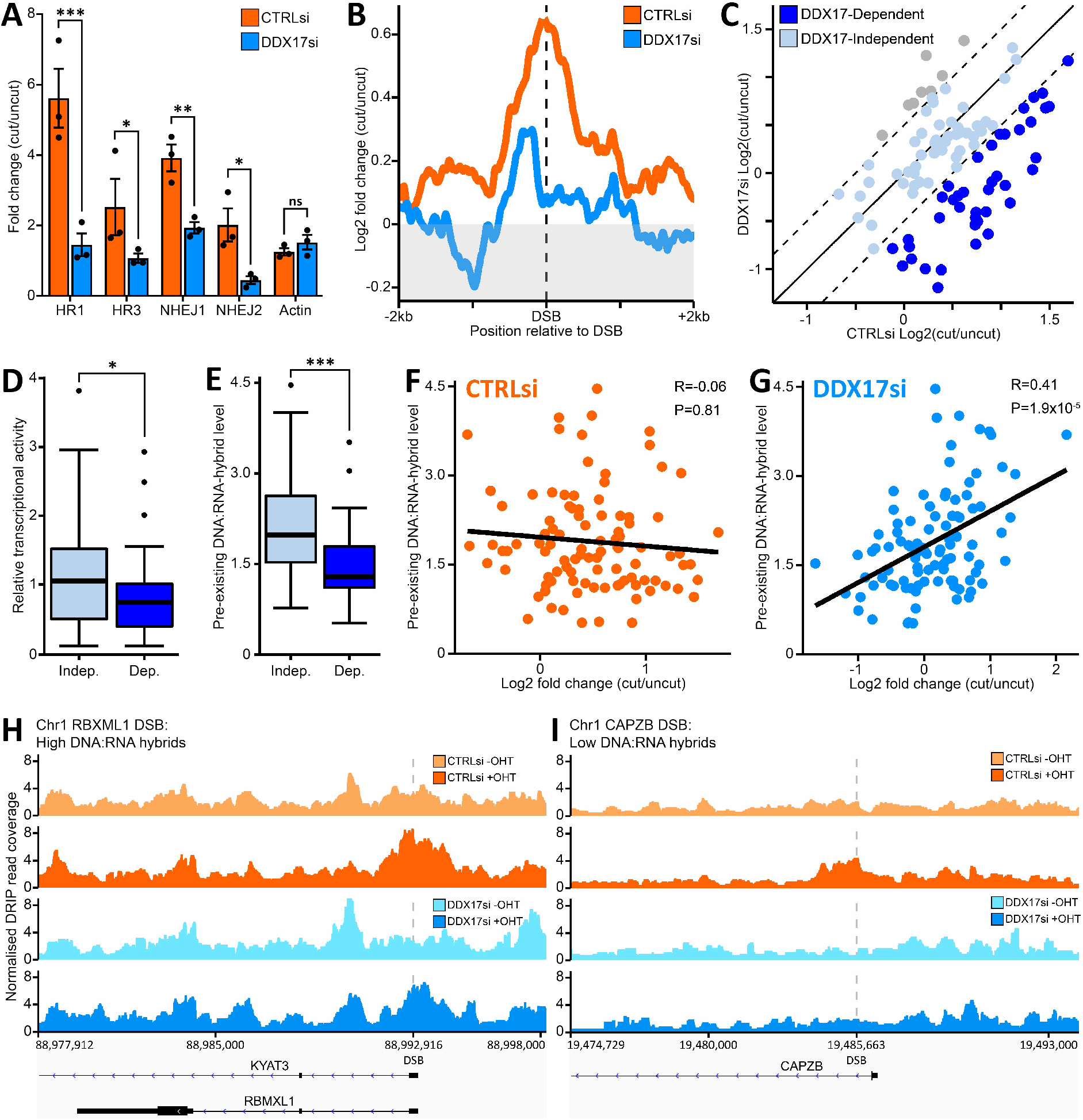
DDX17 promotes double-strand break induced DNA:RNA-hybrids at hybrid deficient genomic loci. (A) DNA:RNA IP (DRIP) qPCR around HR and NHEJ repaired DNA break sites in the DIvA cell system and an undamaged site in actin exon 5. Bar represents the mean fold change of damaged (+OHT) over undamaged (-OHT) for the IP % of input for 3 biological replicates, error bar is SEM. Statistics were done with a paired t-test, *p<0.05, **p<0.01, ***p<0.001. (B) DRIP-seq DSB centred metagene of 99 loci cut by the AsiSI restriction endonuclease in the DIvA cell system, y-axis is log2 fold change of damaged/undamaged normalised readcounts. (C) Log2 fold change of damaged/undamaged DRIP normalised readcounts at 99 AsiSI cut sites in DDX17 siRNA vs control siRNA treatment. Sites close to the y=x line (grey blue) are classed as DDX17-independent, the 40 sites deviating from y=x by less than −0.5 (dark blue) are classed as DDX17-dependent and there are 8 sites (grey) that only slightly deviate from x=y by more than +0.5. (D) Boxplot of relative transcriptional activity of the DDX17-independent DSB loci vs the DDX17-dependent DSB loci, statistics are from directional unpaired Wilcoxon test, *p<0.05. (E) Same as (D) but for pre-existing DNA:RNA-hybrid level, ***p<0.001. (F) Correlation plot of pre-existing DNA:RNA-hybrid level at 99 ASISI induced DSB loci against the Log2 fold change of damaged/undamaged DNA:RNA-hybrid level in the control siRNA treated sample, statistics were done with Pearson correlation testing. (G) Same as (F) but for the DDX17 siRNA treated sample. (H) Genome browser plot of DRIP-seq normalised readcount for −/+ OHT treated with control siRNA and DDX17 siRNA treated samples at the high pre-existing DNA:RNA-hybrid AsiSI induced DSB locus in the RBMXL1 gene. (I) Same as (H) but for the low pre-existing DNA:RNA hybrid level AsiSI induced DSB locus in the CAPZB gene.

As previously reported(12, 17, 24, 56), DRIP-qPCR found that DNA:RNA-hybrid levels increased around the DSB sites in response to induction, but not around an uncut control locus, and that all observed signal was sensitive to RNase-H pre-treatment, indicating that the enrichment is specifically from DNA:RNA-hybrids (Supplementary Figure S4B). Upon DDX17 knockdown, we found that DSB-induced DNA:RNA-hybrids were dramatically reduced at all 4 loci we quantified by qPCR, suggesting that DDX17 is required for the formation of these structures in response to break formation (Fig. 4A, Supplementary Figure S4B). To examine this phenotype at higher resolution, we also conducted high-throughput sequencing of our DRIP samples (i.e. DRIP-Seq), allowing us to analyse the formation of these hybrids across all the sites cut by the AsiSI enzyme in DIvA cells. An overall analysis of the 99 frequently cut AsiSI loci confirmed that DDX17 depletion causes a significant decrease in DSB-induced DNA:RNA-hybrids (Fig. 4B, Supplementary Figure S4C). Comparing AsiSI sites previously reported to be prone to either NHEJ or HR repair(57) found a significant decrease in DSB-induced DNA:RNA-hybrids at both groups upon DDX17 knockdown (Supplementary Figure S4D). In addition, comparing DSB sites with high transcriptional activity to those with low transcriptional activity found that DDX17 again significantly reduced DSB-induced DNA:RNA-hybrids in both groups (Supplementary Figure S4E).

A site-by-site analysis showed that DDX17 knockdown only significantly affects a subset of AsiSI DSB sites; approximately half of these sites show little to no change (Fig. 4C). This allowed us to categorise the sites into either DDX17-dependent, where DDX17 knockdown reduces DSB-induced hybrid formation, or DDX17-independent, where DDX17 knockdown has no significant effect on DSB-induced hybrid formation (Fig. 4C). This classification separates the effects of DDX17 depletion, as the DDX17-dependent sites show no DSB-induced hybrids after DDX17 knockdown whereas the DDX17-independent sites still have a strong induction of DNA:RNA-hybrids levels compared to CTRLsi (Supplementary Figure S4F-H). Classification of these groups allows us to elucidate the factors that contribute to DDX17-dependence of DSB-induced hybrids by comparing the genomic features of these two groups. Interestingly, there were eight sites that showed increased DSB-induced hybrids upon DDX17 knockdown (Fig. 4C), though this increase is quite subtle and there are too few of these sites to accurately investigate this mechanism. We have previously reported that DSB-induced DNA:RNA hybrid formation positively correlates with the pre-existing transcriptional activity of the damaged locus(22). We therefore initially compared the relative transcriptional activity of the DDX17-dependent and independent groups and determined that, although the difference is subtle, DDX17-dependent loci tend to have lower transcriptional activity than the independent sites (Fig. 4D). Since there is a subtle relationship between DDX17-dependency and transcriptional activity, we next chose to investigate features related to transcription. We examined the levels of pre-existing DNA:RNA-hybrid between the groups which showed that DDX17-dependent sites have substantially lower levels of pre-existing DNA:RNA-hybrids than DDX17-independent sites (Fig. 4E) and this differential was proportionally greater than their decreased relative transcriptional activity.

While it is known that transcriptional activity and DNA:RNA-hybrids are strongly linked, they are not directly proportional due to complex regulatory features governing DNA:RNA-hybrids. Recent studies have shown that DNA and RNA-binding proteins, chromatin modifications, chromatin remodelling and processes such as splicing and replication have significant influence on DNA:RNA-hybrid levels(50, 58–63). In the presence of DDX17 we found no correlation between pre-existing DNA:RNA-hybrids and DSB-induced hybrids (Fig. 4F). However, upon DDX17 depletion we see a dramatic shift resulting in a positive correlation between these features (Fig. 4G). Therefore, in the presence of DDX17, DSB-induced hybrids do not rely on pre-existing hybrid levels, but with loss of DDX17 the generation of DSB-induced hybrids becomes dependent on the innate ability of the locus to form these hybrid structures. Interestingly, a re-analysis of our previous DRIP-seq data conducted with *DROSHA* knockdown did not show a significant positive correlation between DSB-induced hybrids and pre-existing hybrids (Supplementary Figure S4I), indicating separate mechanisms for the two proteins. This dependence on DDX17 at sites harbouring low levels of DNA:RNA-hybrids when undamaged can be demonstrated by viewing these features at an individual gene level. For example, when comparing the RBXML1 DSB site which has high pre-existing DNA:RNA hybrids to the CAPZB DSB site which has low pre-existing hybrids (Supplementary Figure S4J). The RBXML1 locus shows DSB-induced DNA:RNA-hybrids in both control and DDX17 knockdowns, whereas the CAPZB locus has little to no induction upon DDX17 knockdown (Fig. 4H-I, Supplementary Figure S4K).

We have determined that DDX17 facilitates DSB-induced DNA:RNA-hybrid formation in the vicinity of DSBs and that this effect is most notable at genomic loci with innately low levels of DNA:RNA-hybrids. In fact, DDX17 depletion leads to DSB-induced hybrid levels being directly proportional to the loci’s pre-existing hybrid levels, suggesting that without DDX17 this DSB-dependent induction of hybrids relies on the loci’s natural ability to form hybrids. As a result, given that they are inefficient at forming hybrids normally, we suggest that DDX17-dependent loci require enzymatic assistance to form DSB-induced DNA:RNA-hybrids, whereas loci of high pre-existing hybrid levels are naturally more efficient at forming DNA:RNA hybrids and are therefore not as dependent upon DDX17.

## DISCUSSION

In this study we used a meta-analysis of 11 published large-scale proteomic datasets investigating DNA repair mechanisms to identify potentially novel DNA repair factors. A number of these publications noted that RNA related gene groups were significantly enriched in their results(40, 42, 44, 64, 65), and some specifically noted DDX17 as a top hit(44, 66). Our analysis yielded a significant number of genes associated with gene expression and splicing, alongside DNA repair genes, as those most commonly identified. In particular, DDX17 was identified with high confidence, as a potential DNA repair gene. Therefore, we chose to investigate the possible role of DDX17 in the DNA damage response.

Clonogenic survival assays revealed that depletion of DDX17 sensitises cells to IR, Olaparib and ICRF-193 indicating a broad role for DDX17 in the response to agents that induce DSBs. Indeed, neutral comet assays found DDX17 depletion to significantly decrease repair of IR-induced DSBs. Consistent with defective DSB repair, metaphase spreads revealed that DDX17 is necessary to prevent chromosomal loss post IR. Thus, DDX17 is important for the maintenance of genome stability.

Localisation of certain DSB repair factors or specific posttranslational modification to focal repair structures demonstrated that while H2AX phosphorylation was not affected by DDX17 depletion, recruitment of both 53BP1 and BRCA1 was significantly reduced. Defective recruitment of these regulators of DSB repair pathway choice suggests defective NHEJ and HR pathways of repair, which was confirmed in NHEJ- and HR-specific reporter assays. Consistent with a function upstream of 53BP1 and BRCA1, recruitment of RNF168 was significantly DDX17-dependent while RNF8 was DDX17-independent, thus placing the function of DDX17 in signalling DNA damage between these two E3 ubiquitin ligases. As 53BP1 and BRCA1 compete for recruitment to the ubiquitylated H2A and H2AX generated by RNF168, a role for DDX17 in regulating RNF168 can also be inferred from DDX17-dependent formation of conjugated ubiquitin and mono-ubiquitylated γ-H2AX at sites of DNA damage. Competition between 53BP1 and BRCA1 recruitment to RNF168-dependent ubiquitylation of histones is a key regulatory point in the choice between HR and NHEJ and subject to complex regulation (10-12, 67-69). It is likely that this stage of the DDR is subject to even more complex regulation than initially believed due to many fine-tuning controls over downstream pathway choice.

DNA:RNA-hybrids are now known to rapidly form at DSBs and are required for efficient repair(12, 17, 19). Furthermore, several RNA binding proteins having been reported to regulate the formation of DNA:RNA-hybrids around DSBs(12, 17, 23, 24, 70). As DDX17 is known to interact with DNA:RNA-hybrids(50), we examined a potential role for DDX17 in regulating DSB-induced DNA:RNA hybrids by quantifying and mapping these hybrids across the genome. We found that DDX17 promotes the formation of DSB-induced DNA:RNA-hybrids and this role is particularly important at loci that normally have low levels of pre-existing DNA:RNA-hybrids. Upon DDX17 depletion, DSB sites in loci with low pre-existing DNA:RNA-hybrids were unable to form the DSB-induced hybrids necessary to propagate an efficient damage signal(12, 20, 71). However, the role of DDX17 in stimulating these hybrids at loci normally expressing high levels of DNA:RNA hybrids was less important as such loci could form sufficient DSB-induced hybrids even upon depletion of DDX17. We hypothesise that loci with naturally low DNA:RNA-hybrid levels require enzymatic assistance from DDX17 to efficiently form DSB-induced hybrids as they lack features, such as preferential DNA sequence and chromatin structure(72, 73), required for high levels of hybrid formation. Since it has been shown previously that these DSB-induced hybrids are necessary for efficient DSB signalling and repair(12, 20, 71), we propose that DDX17’s role in the DDR is to facilitate the formation of DSB-induced hybrids, a role that is particularly important at loci with normally low levels of DNA:RNA hybrids.

Several canonical DNA repair genes have also been shown to function via RNA and DNA:RNA-hybrid related processes. For example, the HR factor RAD52 was found to facilitate RNA-dependent DNA repair via strand-invasion of RNA into double-stranded DNA to form a DNA:RNA-hybrid(20, 74, 75). A previous study also reported that the NHEJ complex can associate with DNA:RNA-hybrid at DSBs and that this was necessary for error-free repair(19). Future studies will be required to determine whether DDX17 has functions that interplay with these known DNA repair genes and DNA:RNA-hybrids. However, as DDR ubiquitylation of histones and signalling is DDX17-dependent, it is likely that DDX17-dependent regulation of DNA:RNA hybrids contributes to the remodelling of chromatin flanking DSBs, which is known to be essential for DSB signalling(76–78). This is consistent with the well described ability of DNA:RNA-hybrids to regulate gene expression and chromatin architecture(79, 80). A role for DNA:RNA hybrids in chromatin remodelling at DSBs would explain how the loss of hybrid promoting proteins such as DDX17 and DROSHA impact upon the early DDR cascade.

Elucidating the role of DSB-induced DNA:RNA hybrids in signalling and repairing DNA damage will require identification and full characterisation of all the proteins regulating these structures. The identification of DDX17 as a new contributor to efficient DNA double strand break repair via both major repair pathways, NHEJ and HR, advances our knowledge of the emerging roles for RNA in DNA-repair.

## AVAILABILITY

DRIP-seq analysis scripts are available at https://github.com/Bushell-lab/drip_seq

## ACCESSION NUMBERS

DRIP-seq fastq files are available at ArrayExpress, accession E-MTAB-10814.

## SUPPLEMENTARY DATA

Supplementary Data are available at NAR online.

## AUTHOR CONTRIBUTIONS

ASB, JL, BRH, GLS and WTL performed experiments and data analysis. ASB performed the proteomic meta-analysis. ASB, JL, NL and MB contributed to experimental design and wrote the manuscript. MB and NFL both coordinated the project.

