## Supplemental file for "DDX17 is required for efficient DSB repair at DNA:RNA hybrid deficient loci"

A

| Study | Group | Target | Damage type | Method |
|---|---|---|---|---|
| Boeing et al., 2016 | Damaged Chromatin Associated | Chromatin associated proteins | UV - 30 J/m$^2$ | SILAC |
| Izhar et al., 2015 | Damaged Chromatin Associated | γ-H2AX colocalization | UV micro-irradiation | Immunofluorescence |
| Maréchal et al., 2013 | Damaged Chromatin Associated | RPA-ssDNA associated | None | In-vitro RPA-ssDNA-Biotin |
| Chou et al., 2010 | Damaged Chromatin Associated | Chromatin associated proteins | UV - 50 J/m$^2$ | SILAC |
| Abbasi et al., 2019 | DDR-Factor Interactors | KU70 Interactome | None | BioID |
| Xing et al., 2015 | DDR-Factor Interactors | NHEJ-complex interactome | None | AP-MS |
| Hill et al., 2014 | DDR-Factor Interactors | BRCA1 interactome | None | TAP-MS |
| Boeing et al., 2016 | Phosphorylated | Phosphoproteomics | UV - 30 J/m$^2$ | SILAC |
| Bennetzen et al., 2010 | Phosphorylated | Phosphoproteomics | IR - 6 Gy | SILAC |
| Matsuoka et al., 2007 | Phosphorylated | Phosphoproteomics | IR - 10 Gy | "pS/pT-Q" IP SILAC |
| Povlsen et al., 2012 | Ubiquitylated | Ubiquitylomics | UV - 40 J/m$^2$ | SILAC |

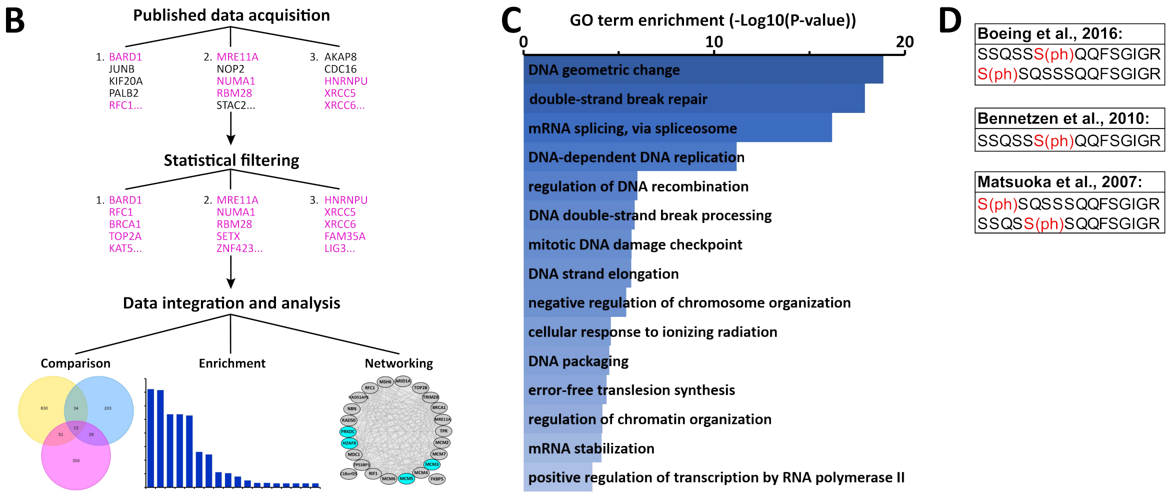


**Supplementary Figure 1**: Design of a meta-analysis of proteomic datasets to identify novel DNA repair factors. **(A)** Table of the 11 studies included in the meta-analysis and information related to them. **(B)** Workflow for the meta-analysis: Results work accessed from publications, filtered for significant results and then integrated into various analyses. **(C)** Gene ontology enrichment analysis of all genes identified in 4 or of the studies in the meta-analysis. **(D)** DDX17 phosphopeptides identified by the three phosphoproteomic studies in the meta-analysis.

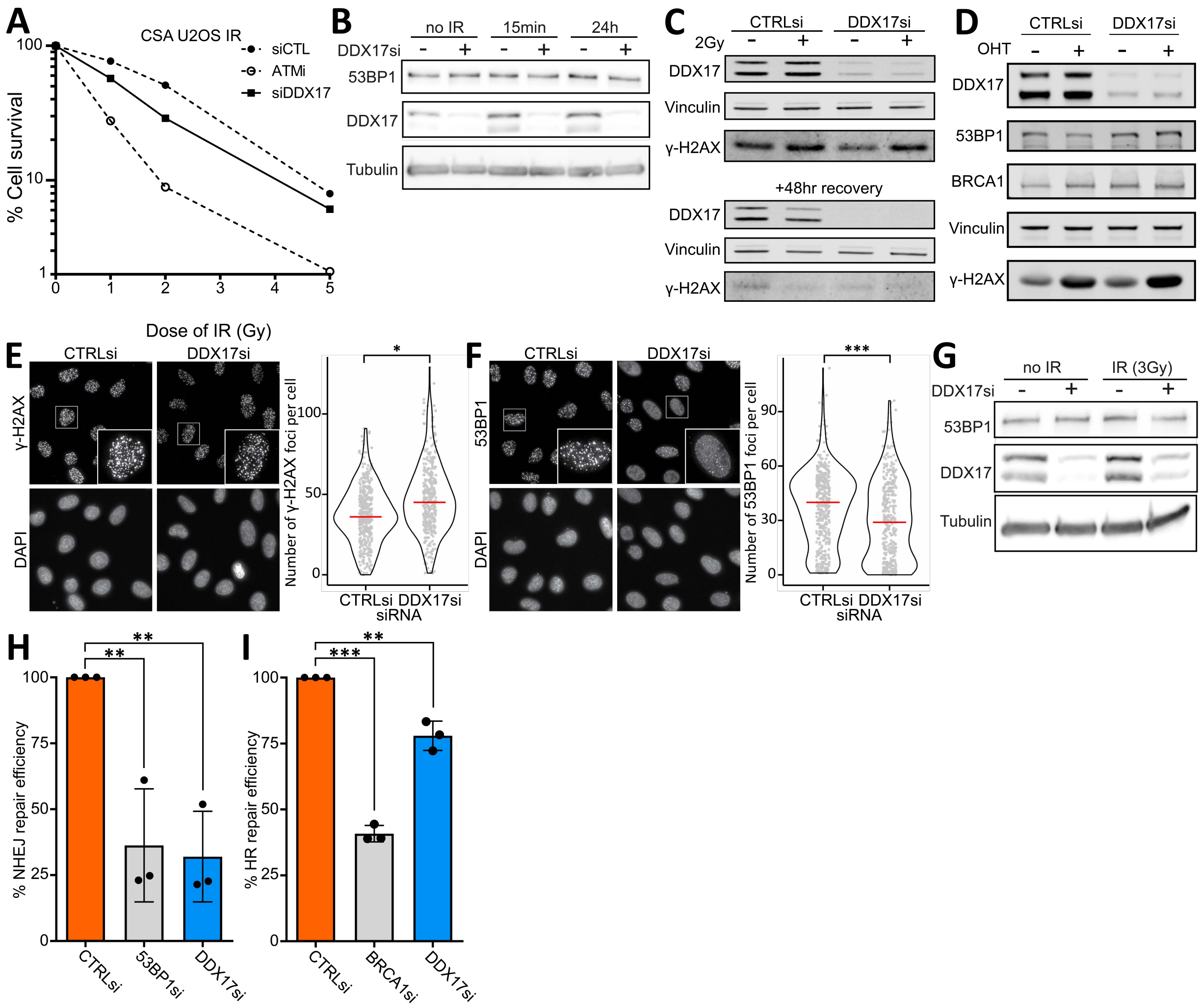


**Supplementary Figure 2**: DDX17 is required for efficient repair of DNA double-strand breaks. **(A)** Clonogenic assay comparing U2OS cell survival in response to increasing doses of IR between the indicated knockdowns and treatments, error bar is SEM across 3 biological replicates. **(B)** Western blot of U2OS cells used in comet assays, treated with either control or DDX17 siRNA and with 0Gy or 3Gy IR followed by 15 minutes or 24 hours. **(C)** Western blot of BJ-5ta used in metaphase spreads, treated with either control or DDX17 siRNA and with 0Gy or 2Gy IR and then allowed to recover for 48 hours. **(D)** Western blots for of DIvA cells transfected with either control or DDX17 siRNA and treated with or without 4-hydroxytamoxifen (OHT). **(E)** Left: representative images of immunofluorescence of γ-H2AX in U2OS cells treated with 3Gy IR with either control or DDX17 siRNA, right: γ-H2AX foci per cell quantification of a minimum of 340 cells per condition across 3 biological replicates, red line is the median. Statistics done using an unpaired directional Wilcoxon test, *p<0.05. **(F)** same as (E) but for 53BP1 immunofluorescence, ***p<0.001. **(G)** Western blot of conditions of (E-F) treated with either control DDX17 siRNA and 0Gy or 3Gy **(H)** EJ5 NHEJ repair reporter assay for 53BP1 and DDX17 knockdowns. Bar height is mean of replicates, points are individual replicate values, error bar is standard deviation, statistics were done using paired t-test, **p<0.01. **(I)** Same as (H) but for the DR-GFP HR repair reporter assay, ***p<0.001.

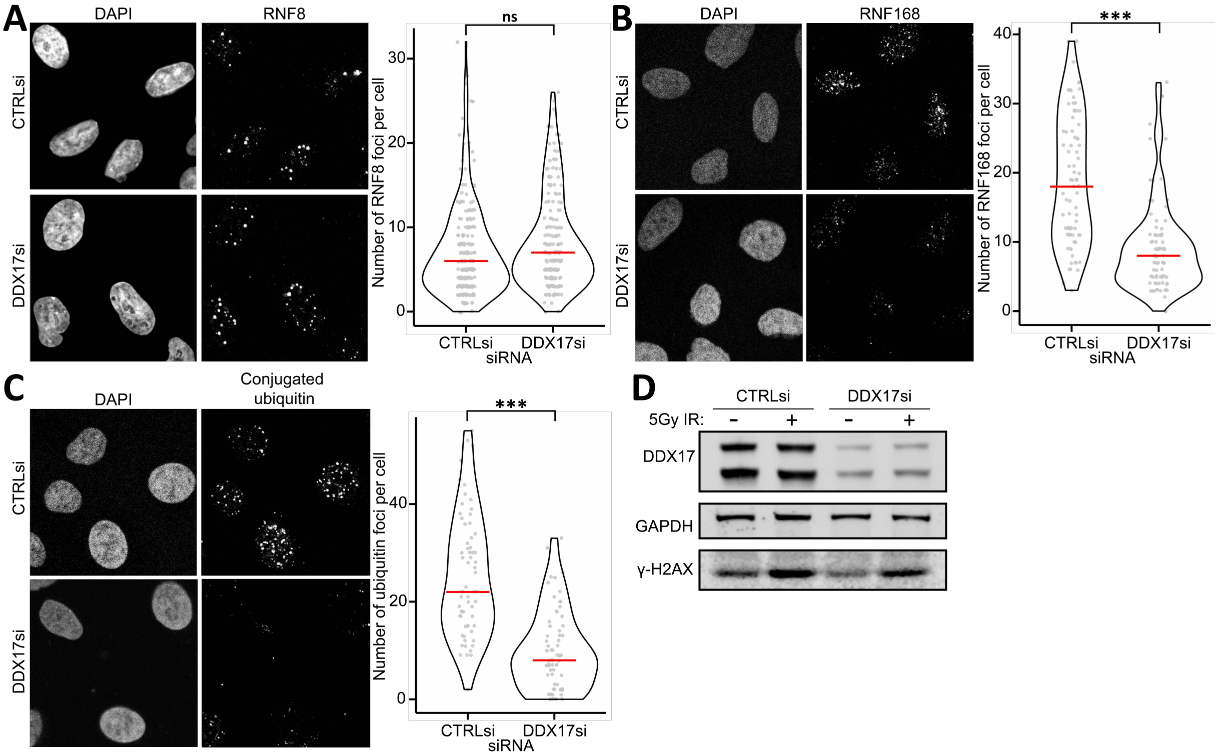


**Supplemental Figure 3**: DDX17 is required for RNF168 recruitment and histone ubiquitylation at DSBs. **(A)** Left: representative images of immunofluorescence of RNF8 in A549 cells treated with 5Gy IR with either control or DDX17 siRNA, right: RNF8 foci per cell quantification of a minimum of 198 cells across 3 biological replicates, red line is the median. Statistics were done using an unpaired directional Wilcoxon test not significant (ns). **(B)** Same as (A) but for RNF168 immunofluorescence in a minimum of 80 cells across 2 biological replicates, ***p<0.001. **(C)** Same as (A) but for conjugated ubiquitin immunofluorescence in a minimum of 68 cells across 2 biological replicates, ***p<0.001. **(D)** Western blot of DDX17 knockdown and 5Gy IR treatment in A549 cells.

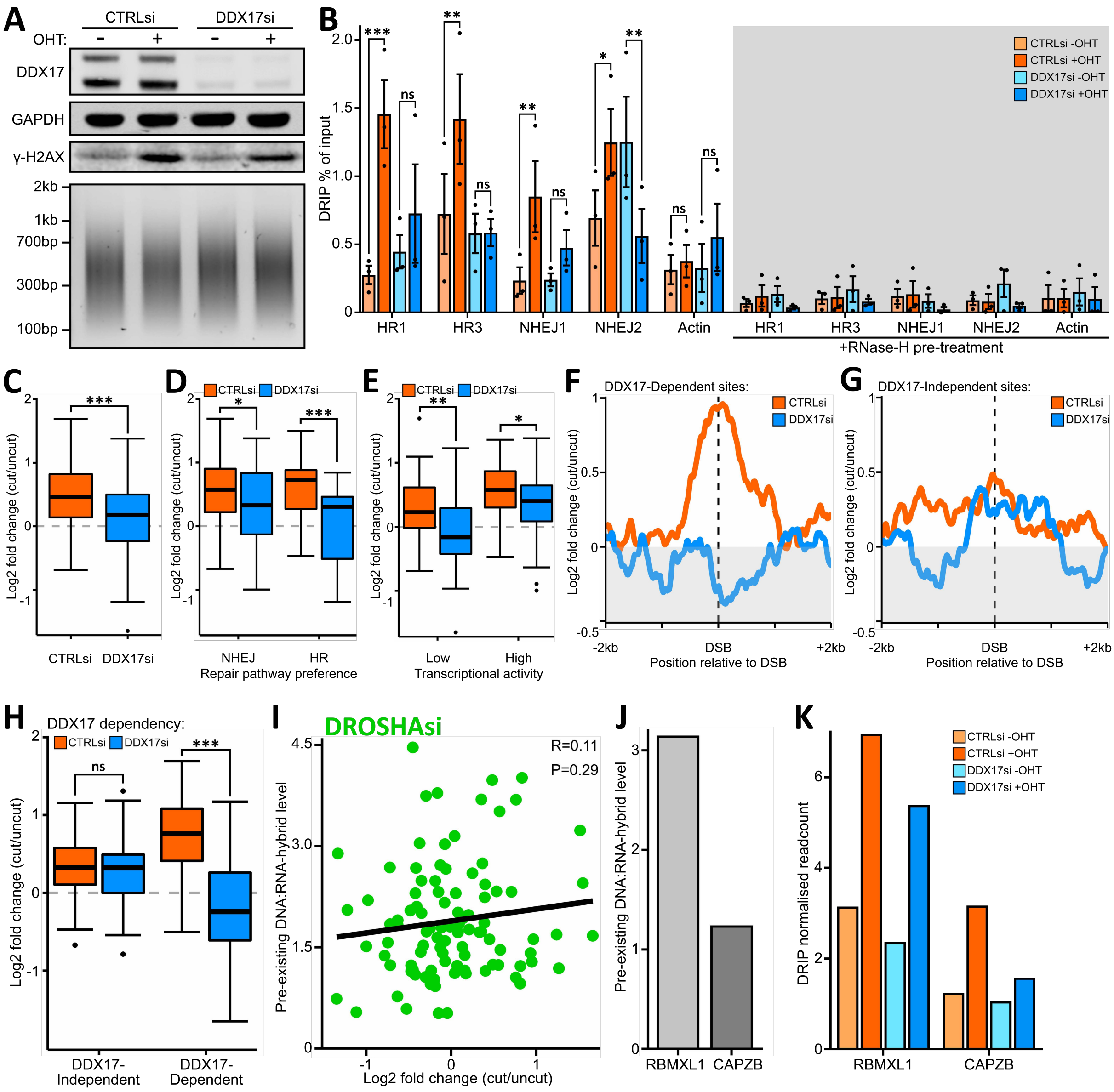


**Supplemental Figure 4**: DDX17 generates DNA:RNA-hybrids around DSBs at low DNA:RNA-hybrid loci. **(A)** Top is western blot validating DDX17 knockdown and 4-hyroxytamoxifen treatment in DIvA cells, bottom is agarose gel validating DNA sonication fragmentation pattern for the DRIP. **(B)** DNA:RNA IP (DRIP) qPCR around HR and NHEJ repaired DNA break sites and an undamaged site in actin exon 5 in DIvA cells treated with 4-hydroxytamoxifen treatment and with either control or DDX17 siRNA. Bar represents the mean % of input for the IP of 3 biological replicates, error bar is SEM, points are individual replicate values. Right is the RNase-H pre-treated negative controls. Statistics were done with a paired t-test, *p<0.05, **p<0.01, ***p<0.001. **(C)** Boxplot of DRIP-seq log2 fold change of damaged/undamaged DRIP read coverage at the ASISI induced DSB sites treated with either control or DDX17 siRNA. Statistics done with an unpaired, directional Wilcoxon test, ***p<0.001. **(D)** Same as (C) but DSB sites are split between those prone to either NHEJ or HR repair, *p<0.05. **(E)** Same as (C) but DSB sites are split between into high and low transcriptional activity groups, **$p<0.01$. **(F)** DRIP-seq DSB centred metagene analysis of DDX17-dependent loci cut by the AsiSI restriction endonuclease in the DIvA cell system, y-axis is log2 fold change of damaged/undamaged normalised readcounts. **(G)** Same as (F) but for DDX17-independent DSB loci. **(H)** Same as (C) but DSB sites are split into DDX17-dependent and DDX17-independent groups. **(I)** Correlation plot of pre-existing DNA:RNA-hybrid level at 99 ASISI induced DSB loci against the Log2 fold change of damaged/undamaged DNA:RNA-hybrid level in DIvA cells treated with *DROSHA* siRNA, statistics were done with Pearson correlation testing. **(J)** Pre-existing DNA:RNA-hybrid level at the AsiSI induced DSB sites in the RBMXL1 and CAPZB genes. **(K)** DRIP read coverage at the AsiSI induced DSB sites in the high DNA:RNA-hybrid RBMXL1 locus and the low DNA:RNA-hybrid CAPZB locus in DIvA cells treated with or without 4-hydroxytamoxifen and with either control of DDX17 siRNA.

**Supplemental Table 1:** List of antibodies used for western blot (WB) or immunofluorescence (IF).

| Antibody Target | Application | Supplier | Catalogue number | Dilution WB | Dilution IF |
|---|---|---|---|---|---|
| DDX17 | WB | SCBT | sc-271112 | 1:1000 | N/A |
| 53BP1 | WB, IF | Novus | NB100-305 | 1:1000 | 1:600 |
| BRCA1 (D-9) | WB, IF | SCBT | sc-6954 | 1:500 | 1:100 |
| Vinculin | WB | Abcam | Ab18058 | 1:5000 | N/A |
| γ-H2AX | WB, IF | Merck | 05-636 | 1:1000 | 1:600 |
| RNF8 | WB, IF | SCBT | sc-271462 | 1:500 | 1:50 |
| RNF168 | WB, IF | SCBT | ABE367 | 1:500 | 1:100 |
| Conjugated Ubiquitin (FK2) | IF | Merck | 04-263 | N/A | 1:200 |
| GAPDH | WB | SCBT | sc-32233 | 1:10,000 | N/A |
| DDX17 | WB | Abcam | ab24601 | 1:1000 | N/A |
| 53BP1 | WB, IF | Novus | NB100-304 | 1:2500 | 1:500 |
| Tubulin (B512) | WB | Sigma | T5168 | 1:10,000 | N/A |

**Supplemental Table 2:** List of primers used in DRIP-qPCR.

| Target | Primer sequences | Amplicon (hg38) |
|---|---|---|
| HR1 | FWD:CCGCCAGAAAGTTTCCTAGA<br>REV:CTCACCCTTGCAGCACTTG | chr22:38468175-38468329 |
| HR2 | FWD:CCTAGCTGAGGTCGGTGCTA<br>REV:GAAGAGTGAGGAGGGGGAGT | chr20:32358116-32358311 |
| NHEJ1 | FWD:ATCGGGCCAATCTCAGAGG<br>REV:GCGACGCTAACGTTAAAGCA | chr6:89638541-89638692 |
| NHEJ2 | FWD:GGTGCCCACAGCTCTCTATG<br>REV:GAAGCCAGAGGAGTGTCCTG | chr9:29212545-29212742 |
| Actin | FWD:GTGACACAGCATCACTAAGG<br>REV:ACAGCACCGTGTTGGCGT | chr17:81511013-81511146 |
